# Liquid-Infused Silicone Catheters Reduce Fungal Burden and Inflammation in *Candidozyma auris* Bladder Infections

**DOI:** 10.64898/2026.02.05.703977

**Authors:** Alyssa Ann La Bella, Hope Akegbe, Caitlin Howell, Felipe H. Santiago-Tirado, Ana L. Flores-Mireles

## Abstract

*Candidozyma auris* is a high-priority, emerging fungal pathogen frequently isolated from urine in healthcare settings. These isolates are often associated with indwelling urinary catheters, a primary risk factor for catheter-associated urinary tract infections (CAUTIs). Despite its clinical prevalence, the mechanisms of *C. auris* colonization and pathogenesis within the bladder remain poorly understood. In this study, we screened *C. auris* isolates from diverse clades using an *in vitro* biofilm model and *in vivo* murine models of uncomplicated UTI and CAUTI. While *in vitro* biofilm formation varied among isolates, the presence of a catheter *in vivo* significantly enhanced fungal burden in the bladder. Notably, one strain (B11103) caused rapid systemic dissemination and mortality. To address this, we evaluated a liquid-infused silicone (LIS)-catheter coating, which has previously shown efficacy against other uropathogens. The LIS-coating significantly reduced *C. auris* attachment *in vitro* and, crucially, mitigated fungal burden on both the catheter and bladder tissue *in vivo* across all tested strains. For the hyper-virulent B11103 strain, LIS-catheters also significantly reduced dissemination to the kidneys and bloodstream. Furthermore, cytokine analysis revealed that *C. auris* CAUTI upregulates IL-6, CSF3, and CXCL1; notably, this damaging inflammatory response was dampened by the LIS-catheter. These findings demonstrate that catheterization potentiates *C. auris* pathogenicity and identify LIS-catheters as a promising, antimicrobial-sparing strategy to prevent colonization, systemic spread, and inflammation during *C. auris* CAUTI.

**IMPORTANCE:** This research addresses the critical public health challenge posed by the emergence of *Candidozyma auris,* elucidating its pathogenesis in the urinary tract, the second most common yet understudied reservoir. Here, we find that *C. auris* exhibits plasticity in its ability to form biofilms in urine and cause uncomplicated UTIs and catheter-associated UTIs. Importantly, we show that our liquid-infused silicone (LIS)-catheters effectively disrupt this cycle by reducing fungal burden, preventing systemic spread, and dampening the damaging host inflammatory response. This work establishes the urinary tract as a critical niche for systemic entry and provides a validated strategy for infection prevention.

## INTRODUCTION

*Candidozyma auris* (formerly *Candida auris*) is a fungal pathogen of critical concern in healthcare settings due to its strong adherence properties, antifungal resistance, and high mortality rates^1–4^. Originally isolated from an ear canal in 2009, *C. auris* is now frequently recovered from the axilla, groin, nares, urine, blood, and skin^4,5^. As an emerging pathogen, our understanding of *C. auris* biology and pathogenesis within these distinct host environments remains limited^3,6,7^. Beyond human hosts, the strong adherence properties of *C. auris* extend to inanimate objects, turning hospital surfaces and medical devices into reservoirs for outbreaks^5–7^.

Urinary catheters are one of the most prevalent medical devices, utilized in up to ∼25% of hospitalized patients and ∼22% in nursing homes, for indications such as urinary retention, anatomic obstruction (e.g. prostatic hypertrophy), surgery, neurogenic bladder dysfunction, or monitoring critically ill patients^8–10^. However, catheterization is the primary driver of catheter-associated urinary tract infections (CAUTIs)^11^. CAUTIs are a leading cause of hospital-acquired infection, with risk approaching 100% for long-term catheter users, and are associated with severe complications, including sepsis and mortality, as well as substantial healthcare costs^11–13^. Despite affecting the bladder, CAUTI pathophysiology differs from uncomplicated urinary tract infections (uUTIs)^11,14^. uUTIs occur in relatively healthy individuals without urinary tract structural abnormalities or immunocompromising conditions, are frequently caused by uropathogenic *E. coli*, and disproportionately affect women^11,14,15^. In contrast, CAUTIs affect patients of both sexes and involve diverse, often multidrug-resistant pathogens, including fungi^14^.

*Candida* spp, specifically *C. albicans*, are the second most prevalent cause of CAUTI^11,16^. Although *C. albicans* rarely causes uUTIs in the dynamic environment of a non-catheterized bladder, the placement of a urinary catheter significantly facilitates infection^11,14,16^. *C. albicans* establishes catheter-associated infections through Efg1-regulated biofilms, which are anchored by the binding of adhesin Als1 to host fibrinogen (Fg) on the catheter and bladder surface^16,17^. Although *C. albicans* drives the majority of fungal CAUTIs, the contribution of *C. auris* to this burden is undefined.

Recent global surveillance of *C. auris* isolation sources indicates that approximately 15% of 12,996 isolates were recovered from urine^5^. By 2024, this proportion rose to nearly 20%, establishing urine as the second most common source of isolation^5^. While it is unclear what proportion of these cases coincided with catheterization, anecdotal evidence suggests a strong association between urinary catheters and multidrug-resistant *C. auris* isolates, a factor that increases mortality risk^5^. Consequently, the capacity of *C. auris* to establish bladder infections in the absence of a catheter remains unknown and requires further investigation.

Given the commonality and severity of CAUTIs, significant effort is currently directed toward developing novel catheter coatings and materials^18–20^. We previously demonstrated that liquid-infused silicone (LIS)-catheters significantly reduced both bladder and catheter microbial burdens in a murine CAUTI model^18^. This efficacy was observed across a broad spectrum of pathogens, including *E. faecalis, E. coli, P. aeruginosa, K. pneumoniae, A. baumannii,* and *C. albicans*^18^. Mechanistically, the LIS coating inhibits the deposition of host proteins, such as Fg, thereby diminishing the scaffold available for pathogen-biofilm formation^18^. Furthermore, as an antimicrobial-sparing therapeutic, LIS-catheters represent a promising strategy for the prevention of CAUTIs and other medical device-associated infections.

Motivated by the frequent isolation of *C. auris* from urine and its threat as a nosocomial pathogen, we investigated its pathogenesis within the urinary tract. We screened a diverse library of *C. auris* isolates from various clades and sources using an *in vitro* CAUTI biofilm model. Subsequently, we evaluated two urine isolates with distinct biofilm phenotypes and one skin isolate using a murine model of uUTI and CAUTI. We observed that while the presence of a urinary catheter significantly enhances fungal burden in the bladder, the use of Liquid-Infused Surface (LIS)-catheters mitigates both bladder and catheter colonization and systemic dissemination. Importantly, while *C. auris* strains induce varying degrees of immune activation, the use of LIS-catheters effectively suppresses both the local bladder inflammation and the severe systemic cytokine response typically seen during CAUTI and with aggressive strains like B11103. Ultimately, these findings demonstrate that LIS-catheters prevent infection by inhibiting fungal attachment and dampening the associated inflammatory response.

## RESULTS

### Urine is a common site of *C. auris* isolation

Previously, we analyzed metadata from 12,996 clinical strains (2004–2024) and identified urine as a common *C. auris* reservoir, ranking second in the US and third globally^21^. To assess prevalence in 2025, we retrieved 8,912 clinical isolates from NCBI’s Pathogen Database, with the US reporting the highest volume (**Fig. 1A**). After excluding 1,810 isolates with unspecified sites and 45 from unidentified swabs, we analyzed the remaining 7,055 strains. Of these, 1,421 (20.4%) were recovered from urine or urinary catheters, maintaining urine as the second most common source worldwide (**Fig. 1B**). In the US specifically, after removing isolates with undefined sources, urine remained the second most common source among the 6,798 isolates (**Fig. 1C**). Overall, the US accounted for the majority of global urinary isolates (**Fig. 1D**).

**Figure 1.**
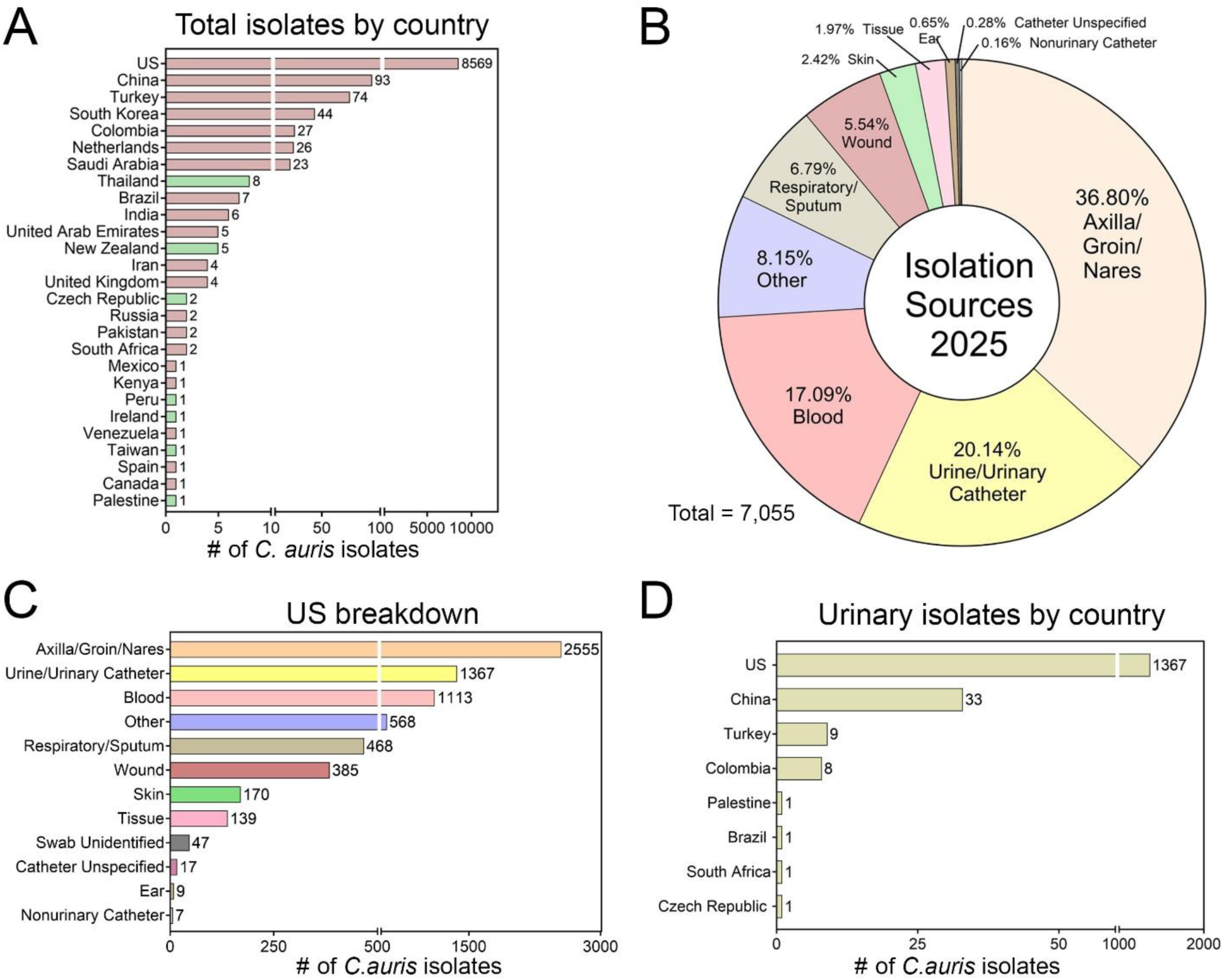
*C. auris* prevalence and isolation sources in 2025. **A**) The number of *C. auris* isolates in 2025 by country according to the NCBI Pathogen Database. Green bars indicate countries that did not previously have *C. auris* isolates from 2004-2024. **B**) Composition of *C. auris* isolates worldwide in 2025. **C**) Breakdown of the isolation sources of *C. auris* in the US. (**D**) Urinary isolates by country in 2025.

### Diverse *C. auris* isolates can form biofilms in CAUTI *in vitro* conditions

Given its prevalence in the urinary tract and on catheters, we evaluated the pathogenic potential of *C. auris* as a CAUTI pathogen. To do this, we assessed biofilm formation using our previously described *in vitro* Fg-urine model, which mimics the catheterized bladder environment^16^. A library of isolates representing Clades I through V and diverse isolation sources (wound, blood, ear, skin, respiratory, nose, groin, or urine) revealed significant variation in biofilm phenotype (**Fig. 2A**). Clade I and IV displayed greatest heterogeneity, containing both high and low biofilm formers (**Fig. 2A**). In contrast, Clade II isolates were consistently low biofilm formers, whereas Clade III isolates produced robust biofilms (**Fig. 2A**). The single Clade V representative also exhibited a high biofilm-forming phenotype (**Fig. 2A**). Importantly, we observed no correlation between isolation source and biofilm capacity; both high and low formers were present across all isolation groups (**Fig. 2A**). This was exemplified by two urine isolates, B11103 (Clade I) and AR1105 (Clade IV), which displayed opposing phenotypes. To better understand if biofilm capacity is due to isolation source or clade, we moved forward with our study characterizing the two urine isolates from different clades (B11103 and AR1105) as well as a skin isolate, Chicago 4, from Clade IV (a gift from Dr. Teresa O’Meara). Fluorescence microscopy confirmed our quantitative biofilm results of these strains, visualizing robust biofilm architecture in AR1105 and the skin isolate Chicago 4, contrasted by a lack of biofilm formation in B11103 (**Fig. 2B**).

**Figure 2.**
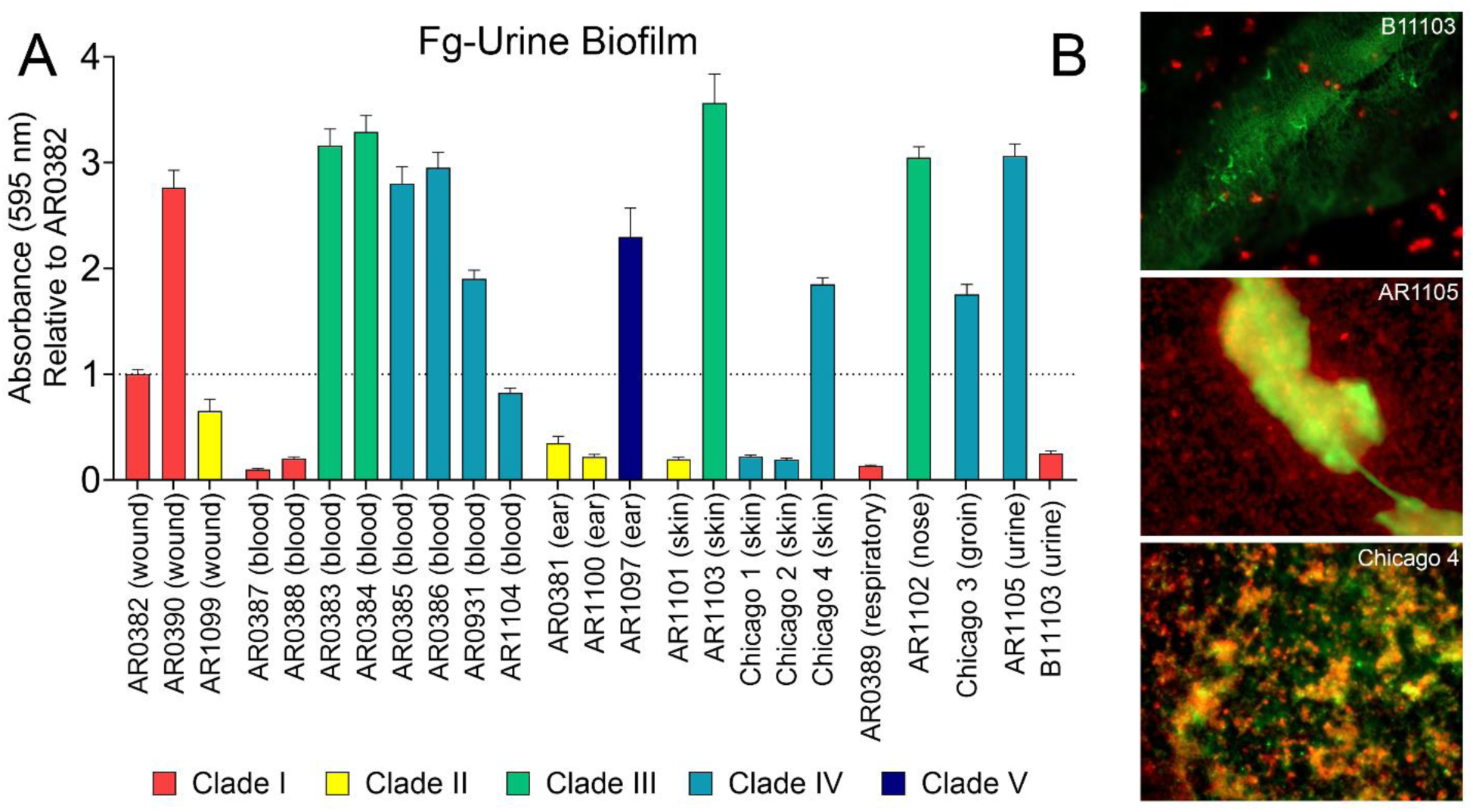
Screening of *C. auris* isolates from different body sites and clades with an *in vitro* catheterized bladder model. **(A)** Biofilm formation of *C. auris* isolates on Fg-coated plastic in urine conditions quantified by crystal violet staining (relative to AR0382). **(B)** Fluorescent microscopy of fibrin(ogen)-urine biofilms at 40x magnification for three *C. auris* isolates: B11103 – Clade I – urine isolate; AR1105 – Clade IV – urine isolate; Chicago 4 – Clade 4 – skin isolate. (*C. auris* red; fibrin(ogen) green).

### Urinary catheterization significantly enhances *C. auris* bladder burden *in vivo*

To determine the uropathogenic potential of *C. auris*, we assessed three isolates (B11103, AR1105, and Chicago 4) at 1-day post-infection (dpi) in established murine models of uUTI and CAUTI^22^. For all three strains, the presence of a urinary catheter significantly increased fungal burden in the bladder (**Fig. 3A-C**). While all isolates successfully colonized the catheter in the CAUTI model, B11103 exhibited exceptional virulence, causing rapid dissemination and mortality, indicated by the hexagon data points (**Fig. 3A**). Notably, we have not observed mortality within 24 hours (<1 dpi) with other urinary pathogens, highlighting the exceptional pathogenic potential of *C. auris* in the catheterized bladder. This hyper-virulent phenotype extended to the uUTI model, where B11103 caused significantly higher bladder burdens and renal dissemination than either AR1105 or Chicago 4 **(Fig. 3A-C**).

**Figure 3.**
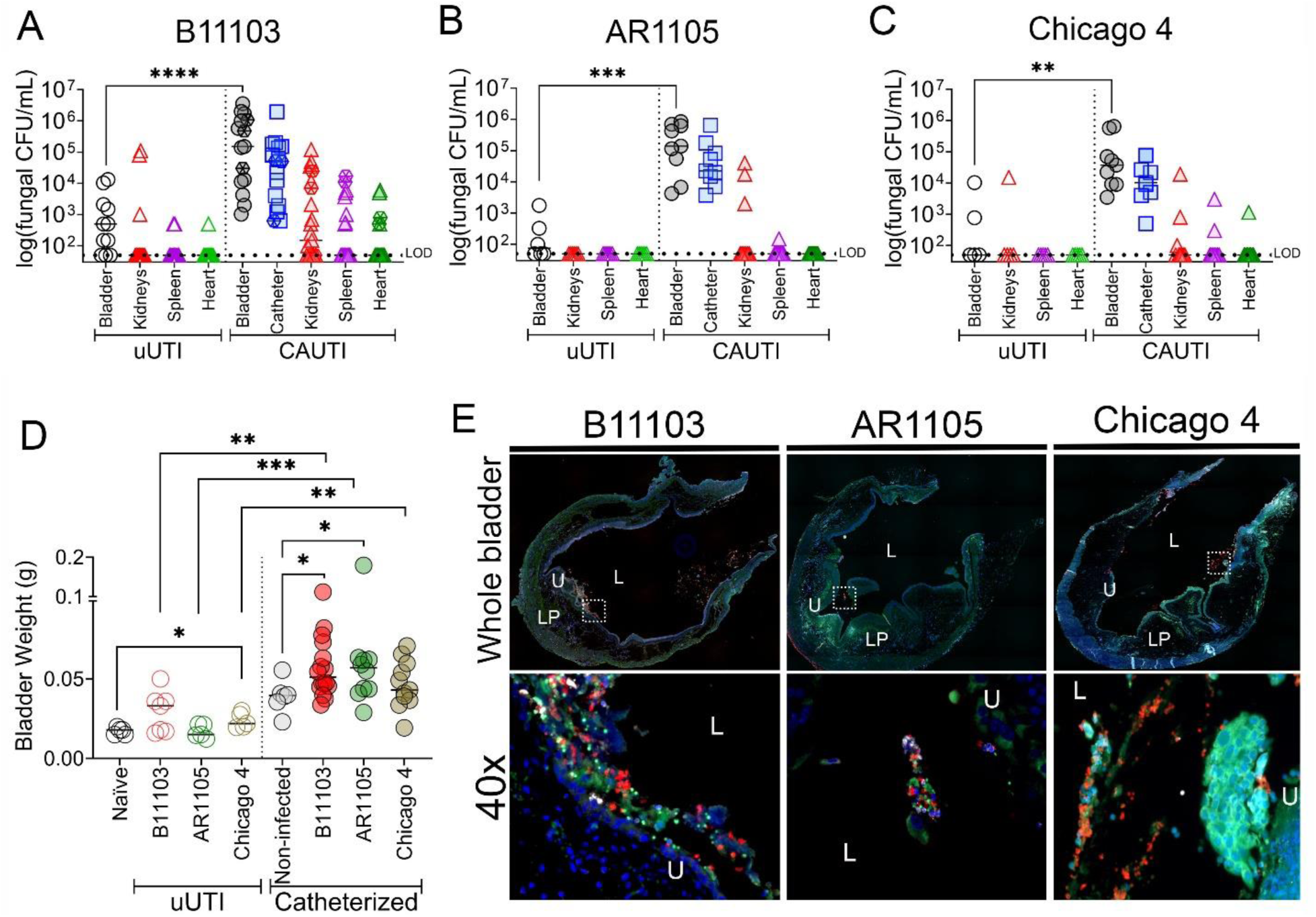
Characterization of *C. auris* pathogenesis in murine models of UTI and CAUTI. Fungal burden of harvested organs in a uUTI and CAUTI mouse model when transurethrally infected with **(A)** B11103, **(B)** AR1105, and **(C)** Chicago 4 after 1 dpi. **(D)** Weights of infected (uUTI) only and catheterized and infected (CAUTI) bladders. **(E)** Immunofluorescence staining of catheterized and infected bladders after 24 hours (DAPI blue; *C. auris* in red; fibrin(ogen) in green; Ly6G in white) at 10x and 40x magnification. White box indicates the 40x zoom-in. L lumen; U urothelium; LP lamina propria. Differences between groups were tested for significance using the Mann-Whitney *U* test. \**P* < 0.05, \*\**P* < 0.005, \*\*\**P* < 0.0005, and \*\*\*\**P* < 0.0001.

As an initial assessment of inflammation in our uUTI and CAUTI models, we used bladder weight as a proxy (**Fig. 3D**). Compared to naïve (uninfected) controls, uUTI infection increased bladder weights for both strains. This elevation was statistically significant for Chicago 4, while B11103 trended toward significance (**Fig. 3D**). Urinary catheterization is known to cause physical damage; therefore, to understand the impact of the pathogens, we compared the bladder weights to catheterized non-infected controls. We found the two urinary isolates, B11103 and AR1105, had significantly increased bladder weights compared to the catheterized non-infected bladders (**Fig. 3D**). Given the high fungal burden associated with catheterization, we visualized colonization via immunofluorescence (**Fig. 3E**). Imaging revealed all three strains localized throughout the lumen and lining the urothelium, appearing primarily as single yeast cells or small aggregates that were associated with Fg (**Fig. 3E**).

### Liquid infused silicone catheters reduce *C. auris* attachment in urine *in vitro* and mitigates CAUTI *in vivo*

Our previous work demonstrated that liquid-infused silicone (LIS)-catheters effectively reduce uropathogen attachment, bladder inflammation, and protein fouling, specifically Fg deposition^18^. Given that *C. auris* colonization is driven by a high affinity for abiotic surfaces^6,7^ and co-localization with Fg in the catheterized bladder (Fig. 3E), we hypothesized that LIS-catheters would be highly effective against this pathogen. To test this, we quantified adherent fungal cells on unmodified (UM) and LIS-catheter segments in our Fg-urine model. After 48 hours, the LIS-coating significantly reduced biofilm formation across all three *C. auris* isolates (**Fig. 4A**). Importantly, planktonic population remained unchanged (**Fig. 4B**). This dissociation between surface adherence and planktonic growth indicates that LIS-catheters function as a passive, anti-adhesive barrier rather than through biocidal activity.

**Figure 4.**
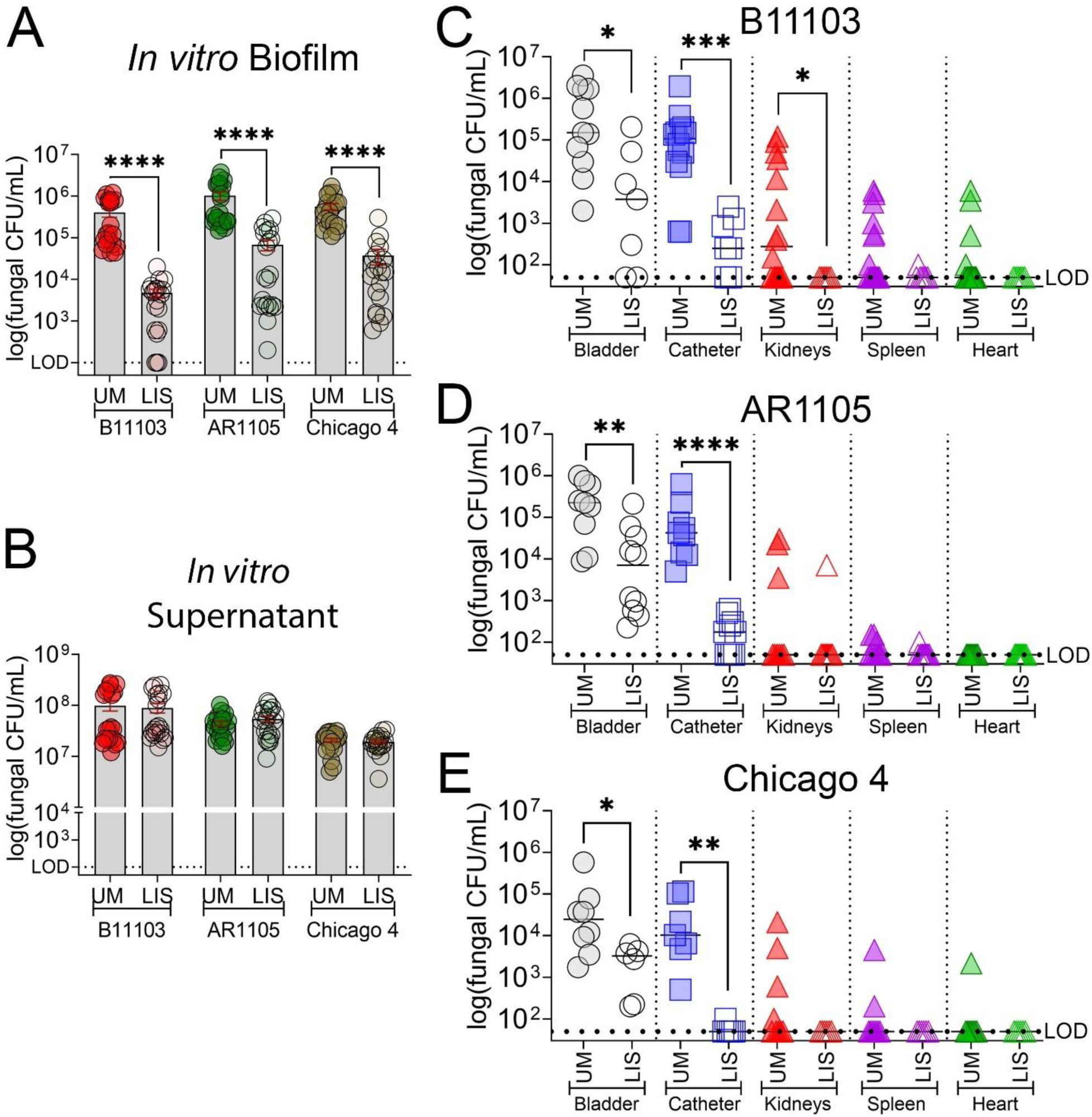
Liquid infused silicone reduces *C. auris* biofilm formation *in vitro* and mitigates infection severity and urosepsis in a mouse CAUTI model. **A**) *In vitro* biofilm formation on unmodified (UM) and liquid-infused silicone (LIS) catheter segments incubated in Fg-urine for 48 hours. **B)** Quantification of fungal cells in urine supernatant of UM and LIS catheters after 48 hours. **C-E)** Comparison of fungal burden in the bladder, catheter, kidneys, spleen, and heart at 24 hours post-infection. Mice were implanted with UM or LIS catheters and infected with (**C**) B11103, (**D**) AR1105, or (**E**) Chicago 4. Differences between groups were tested for significance using the Mann-Whitney *U* test. \**P* < 0.05, \*\**P* < 0.005, \*\*\**P* < 0.0005, and \*\*\*\**P* < 0.0001.

To validate our *in vitro* findings, we assessed the efficacy of LIS-catheters *in vivo*. In our mouse CAUTI model, LIS-catheters significantly reduced fungal burdens in the bladder and on the catheter surface for all strains (**Fig. 4C-E**). Notably, catheter colonization was nearly eradicated in the skin isolate, Chicago 4 (**Fig. 4E**). Furthermore, LIS catheters prevented systemic dissemination, with a statistically significant reduction in kidney colonization observed specifically for the B11103 strain (**Fig. 4C**). Thus, LIS-catheters prove effective at controlling *C. auris* CAUTI and mitigate the risk of ascending infection and potential urosepsis across diverse clades and isolation sources.

### Differential inflammatory response during *C. auris* uUTI and CAUTI

Current knowledge regarding *C. auris* immunopathology is largely limited to skin and bloodstream infection models^23,24^. To define the local response in the urinary tract, we assessed the levels of 23 cytokines in bladder tissue harvested from uUTI and CAUTI mice at 1 dpi. During uUTI, cytokine levels were calculated by comparing fold change over naïve (non-infected and non-catheterized) bladders, finding that B11103 and Chicago 4 induced a higher cytokine response than AR1105 (**Fig. 5A**). This correlates with the higher bladder weights (**Fig. 3D**). Principal component analysis (PCA) revealed two distinct clusters of biological states (non-infected and infected) based on cytokine signatures (**Fig. 5B**). AR1105 elicited a mild inflammatory response, whereas bladders infected with B11103 and Chicago 4 exhibited significantly higher levels of cytokines, including CCL2 (MCP-1), CXCL1 (KC), IL-4, IL-10, and IL-13 (**Fig. 5A-B**).

**Figure 5.**
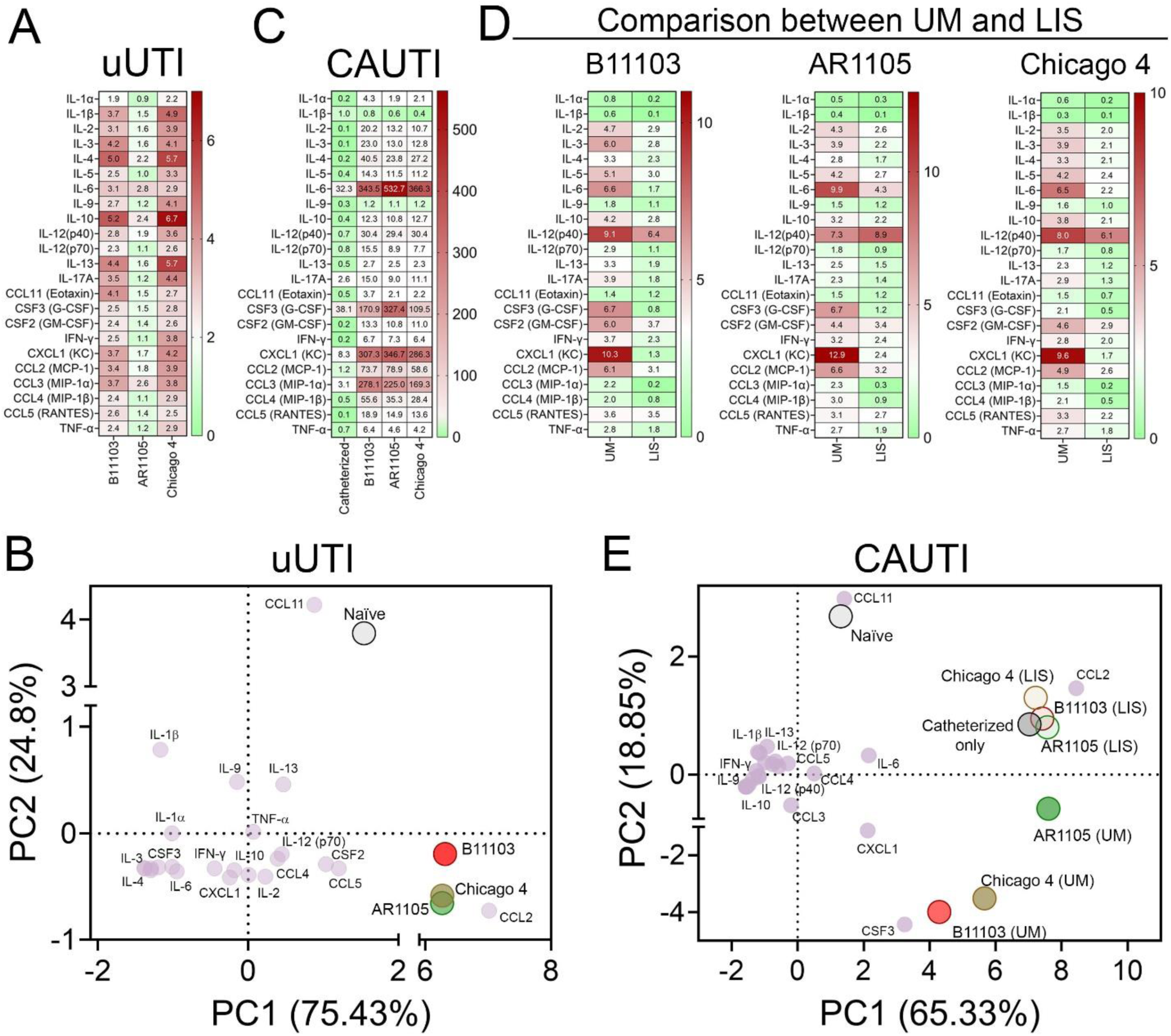
Host-immune response during uUTI and CAUTI. **A)** Bladder cytokine profiles (fold change relative to naïve controls) for mice with uUTI by strains B11103, AR1105, and Chicago 4. **B)** Principal component analysis (PCA) biplot comparing the cytokine signatures of these uUTI groups. **C)** Comparison of cytokine profiles in catheterized-only mice and mice infected and catheterized with UM catheters. Data are normalized to naïve bladder controls to isolate the effect of catheterization and infection. **D**) Bladder cytokine profiles of mice infected with B11103, AR1105, or Chicago 4 and catheterized with UM or LIS catheters. Data were normalized to the catheterized-only control group to compare the efficacy of UM versus LIS catheters. **E)** PCA biplot comparing cytokine signatures of naïve and catheterized-only controls against mice infected with strains B11103, AR1105, or Chicago 4 (UM vs. LIS).

Consistent with previous reports that urinary catheterization alone elicits a strong inflammatory response^13,18,25–27^, we observed elevated levels of IL-6, CSF3 (G-CSF), and CXCL1 (KC) in catheterized controls compared to naïve bladders. Importantly, during CAUTI, all three strains induced a profile dominated by neutrophil-recruiting signals (including CXCL1, IL-6, CSF3, and CSF2) and monocyte/macrophage-recruiting signals (including CCL2, CCL3, and CCL4) (**Fig. 5C**), correlating with the observed bladder inflammation (**Fig. 3D**). We assessed pathogen-driven responses by calculating cytokine fold changes relative to uninfected catheterized controls. Comparative analysis of UM and LIS groups revealed that LIS-catheters significantly reduced inflammatory cytokine levels, even in bladders infected with the highly virulent strain B11103 (**Fig. 5D**). In contrast to other cytokines, IL-12 (p40) levels remained similar to the UM-infected bladders. Consistent with these findings, PCA showed that all UM-infected groups clustered tightly together, characterized by a high-inflammation state driven by pro-inflammatory markers such as CXCL1, IL-6, and CSF3 (**Fig. 5E**). Importantly, LIS-catheterized infected bladders shift significantly away from the high-inflammation UM-infected bladders’ cluster (**Fig. 5D-E**). Their cytokine signatures closely resembled those of uninfected catheterized controls (**Fig. 5E**). These findings correlate directly with the LIS-mediated reduction in fungal burden (**Fig. 4D-E**), indicating that the LIS-coating successfully prevents the infection-induced cytokine storm. In conclusion, LIS technology effectively suppresses inflammation across all *C. auris* tested strains in the CAUTI model.

Given the high systemic dissemination of B11103 observed during catheterization, we analyzed serum cytokine profiles to compare systemic inflammatory responses across strains and between UM– and LIS-catheterized mice. The cytokine data was normalized against serum from catheterized non-infected mice. We found that B11103 elicited the most severe systemic response (**Supplementary Fig. 2A**), correlating with the high dissemination rate (**Fig. 3A**). The cytokine signature was driven by a massive elevation of CSF3, IL-6, IL-1 α, and IL-12(p40) (**Supplementary Fig. 2A**). While Chicago 4 and AR1105 also strongly induced CSF3 and IL-12(p40), IL-6 and IL-1α levels were lower than those of B11103 (**Supplementary Fig. 2A-C**). Importantly, LIS-catheterized and infected mice showed a consistent reduction in circulating cytokines, specifically reducing IL-6 and CSF3 (**Supplementary Fig. 2A-C**). PCA confirmed a differential serum cytokine signature between UM– and LIS-catheterized and infected mice (**Supplementary Fig. 2D**). PCA revealed that LIS-catheterized and infected mice shifted toward a distinct immune signature defined by sustained IL-12(p40) and IL-1α (**Supplementary Fig. 2D**), rather than a neutrophil-recruiting profile (CSF-3 and IL-6) seen in UM-catheterized and infected mice (**Supplementary Fig. 2A-D**). These data demonstrate that LIS technology mitigates strong systemic inflammation driven by CSF3 and IL-6, possibly limiting the local infection burden in the bladder (**Fig. 3 and 4**).

## DISCUSSION

*C. auris’* emergence has created a nightmare for healthcare systems. With little knowledge about its basic biology and pathogenesis, researchers have only just begun to elucidate the fungal pathogen’s behavior in various niches. Recent studies have focused on *C. auris*’ skin colonization as well as characterizing bloodstream isolates^3,21,24,28–30^. In recent years, urine has become the second most common isolation source for *C. auris* (**Fig. 1**)^5^; despite the prevalence, there is a critical lack of studies investigating the fungus within the urinary tract environment.

Here, we screened a library of *C. auris* isolates from varying clades and isolation sources in our *in vitro* biofilm model, observing diverse biofilm phenotypes (**Fig. 2**). Our results suggest that biofilm formation is driven by genetic lineage (clade) rather than clinical isolation source. Specifically, Clade III isolates exhibited robust biofilm potential, contrasting with the low activity of Clade II and the heterogeneity of Clades I and IV. Although confirming these trends requires analysis of more clinical isolates from all six clades (particularly Clade VI, for which isolates were unavailable for this study), the observed lineage-dependency aligns with the pathogen’s genomic landscape. *C. auris* is characterized by significant genetic diversity, defined by at least six highly divergent lineages. While inter-clade differences involve thousands of single nucleotide polymorphism (SNPs), intra-clade diversity arises from smaller variations such as chromosomal rearrangements and aneuploidy, driving diverse resistance and virulence profiles^31^.

This lineage-dependent pathogenicity mirrors patterns seen in other uropathogens, such as uropathogenic *E. coli* (UPEC). UPEC exhibits significant genetic diversity, with specific phylogenetic groups (e.g., B2 and D) carrying distinct virulence factors, including fimbriae and toxins^32–37^. Yet, despite this extensive genetic diversity, UPEC strains exhibit a remarkably conserved gene expression pattern during both human and murine infections^33,36^. Therefore, it is critical to analyze the *in vivo* transcriptional profiles of diverse *C. auris* strains to determine if a similarly conserved transcriptional signature exists during uUTI and CAUTI.

A notable finding in this study is the hyper-virulence of the Clade I isolate B11103 (**Fig. 3**). While virulence heterogeneity is characteristic of *C. auris*, the rapid mortality observed in our CAUTI model represents a distinct departure from the pathophysiology of other typical uropathogens, including *C. albicans*, where acute lethality in murine models has not been observed^16^. Furthermore, we observed a striking inverse relationship between *in vitro* biofilm formation and *in vivo* virulence phenotypes. Specifically, B11103 was a poor biofilm former *in vitro* (**Fig. 2**), yet exhibited high colonization in the catheterized bladder, extensive systemic dissemination, and high lethality *in vivo* (**Fig. 3A**). Bladder imaging revealed *C. auris* yeast cells are able to colonize the urothelium and form biofilms with Fg in the catheterized bladder (**Fig. 3E**). This discrepancy may be explained by several factors. First, nutritional availability differs significantly; the static *in vitro* model lacks fluid replenishment, potentially limiting fungal proliferation, whereas the catheterized bladder constantly provides fresh urine and serum proteins. Alternatively, biofilm dispersion may play a role. The biofilm might be dynamic, undergoing cycles of attachment and detachment, rather than remaining static. This phenomenon parallels *Proteus mirabilis*, where nutrient limitation induces biofilm dispersion to facilitate spreading^38,39^. Indeed, similar temporal dynamics have been observed in *P. mirabilis* Fg-dependent urine biofilms^40^.

Another potential explanation involves an attachment trade-off, a phenomenon well-documented in UPEC. In UPEC, the type 1 pilus, specifically the FimH tip adhesin, is critical for successful bladder infection. FimH binds to mannosylated proteins, including uroplakins on urothelial cells and Fg^14^. Crucially, successful infection requires FimH to function as a ‘catch-bond’ (becoming adhesive only under shear stress from urine flow) rather than acting as a static ‘super-glue’^41,42^. Analysis of FimH sequences has identified natural mutations that lock the adhesin into a permanent high-affinity state; interestingly, these mutations destroy the catch-bond flexibility^41–43^. Consequently, although these mutations confer high mannose binding, they paradoxically lead to reduced UPEC infection capacity^41,42^. This trade-off parallels the behavior of our high biofilm formers, AR1105 and Chicago 4 (**Fig. 2**). It suggests that specific mutations may lock adhesins into static conformations; while this increases attachment, it impairs the dynamic detachment-reattachment cycles required for infecting neighboring cells, evading the immune system, and disseminating systemically.

*C. auris* strains have been famously described as ‘barnacles’ due to specialized adhesion proteins that facilitate strong attachment to human skin, intravenous catheters, and environmental surfaces^7^. However, this raises the question of whether urinary *C. auris* strains are undergoing pathoadaptation, whereby excessive adhesiveness becomes a liability. Instead, a dynamic attachment strategy likely favors immune evasion and systemic dissemination, strategies that are particularly vital in the catheterized bladder since this environment is defined by intense immune surveillance^13,16,25,27^. It would be critical to analyze a broad spectrum of urinary clinical strains to identify if this pathoadaptation is happening. This is clinically critical, as several studies indicate that patients with *C. auris* candiduria (including those with indwelling urinary catheters) experience worse outcomes, often resulting in mortality^44–46^. Therefore, future studies must investigate why certain *C. auris* strains act as localized colonizers while others cause lethal systemic infection during CAUTI.

Among the three isolates tested, B11103 demonstrated superior bladder colonization in the uUTI model. In the absence of a urinary catheter, most CAUTI-associated pathogens are rapidly cleared from the dynamic bladder environment by the hydrodynamic forces of micturition, limited nutrients, and the immune response^47,48^. Although B11103 colonization did not reach the clinical threshold for uUTI (>10^5^ CFUs), it was significantly higher than that of AR1105 and Chicago 4 at 1dpi. (**Fig. 3**). This persistence implies that certain *C. auris* strains possess intrinsic uropathogenic traits similar to UPEC, independent of catheterization. Future studies using low-dose and temporal models are essential to define the extent of this pathogenicity.

Importantly, the catheterized environment promoted colonization of a skin isolate, Chicago IV (**Fig. 3**). This together with the ability of isolates from diverse sources (skin, blood, urine) to form biofilms in our Fg-urine model suggests that *C. auris* is a generalist pathogen with high plasticity, capable of adapting to the urinary niche regardless of its isolation origin. This indicates that the catheter acts as a permissive niche, allowing strains to thrive regardless of their isolation origin. This phenomenon parallels findings in *Enterococcus* species, where diverse lineages (including commensal gut strains) are capable of colonizing the catheterized bladder^49^.

Remarkably, our LIS-catheters significantly inhibited fungal attachment *in vitro* and *in vivo* (**Fig. 4**). In the murine CAUTI model, these modified catheters drastically reduced both bladder burden and catheter colonization across all three tested strains. Notably, for isolate B11103, the LIS catheter effectively mitigated systemic dissemination, significantly lowering fungal loads in the kidneys, spleen, and bloodstream (**Fig. 4**). LIS-catheters also reduced the exacerbated inflammatory response in the bladder, mitigating a cytokine storm characterized by neutrophil and monocyte/macrophage-recruiting signals (**Fig. 5**). Importantly, LIS-catheters decreased the damaging response without compromising IL-12 (p40) levels in the bladder or IL-12 (p40) and IL-1α levels in the bloodstream. These cytokines have been shown to provide critical, non-redundant protection against systemic fungal infections by orchestrating innate and adaptive immune responses. IL-12 drives Th1-cell differentiation and IFN-γ production, enhancing macrophage antifungal activity^50,51^, while IL-1α acts as an “alarmin” to mediate inflammation, reduce tissue damage, and control pathogen growth, particularly in the brain and kidneys^52,53^. Our LIS-catheters significantly inhibited *C. auris* colonization and systemic dissemination, mitigating harmful inflammation while uniquely preserving protective responses. Future studies must now assess the long-term durability of this technology to validate its clinical utility in preventing catheter-associated pathogenesis.

With urine emerging as the second most common site of *C. auris* isolation, our results establish the catheterized urinary tract as a critical yet overlooked reservoir for pathogenesis. These findings indicate that the urinary catheter acts as a permissive gateway, allowing *C. auris* to establish infection regardless of its isolation source. Crucially, the severe and lethal infection caused by the urinary isolate B11103 highlights the urgent need to determine if urinary strains are undergoing pathoadaptation. Given the highly adhesive nature of *C. auris* and its propensity to colonize medical devices, preventing initial attachment is a critical strategy for reducing infection outbreaks. Our validation of LIS-catheters offers a robust countermeasure by inhibiting catheter colonization and subsequent systemic dissemination while simultaneously mitigating the ‘cytokine storm’ without compromising protective IL-12 and IL-1α responses. LIS technology addresses both the physical and immunological challenges of CAUTI. Ultimately, this work redefines urinary *C. auris* not merely as a localized colonizer, but as a significant threat for systemic disease, positioning surface-modified catheters as an essential preventative strategy in clinical settings.

## MATERIALS & METHODS

### Ethics statement

All animal care was consistent with the Guide for the Care and Use of Laboratory Animals from the National Research Council. The University of Notre Dame Institutional Animal Care and Use Committee approved all mouse infections and procedures as part of protocol number 25-01-9009. For urine and blood collections, all donors signed an informed consent form, and protocols were approved by the Institutional Review Board of the University of Notre Dame under study #19-04-5273 for urine and #18-08-4834 for blood.

### Isolation Source Analysis

Using NCBI’s Pathogen Database, we analyzed the clinical isolates of *C. auris* and classified their isolation source into one the following 12 categories: axilla/groin/nares; urine/urinary catheter (urinary); blood; skin; wound; respiratory tract/sputum; ear; tissue (i.e., various bones and tissue types); unspecified catheter; non-urinary catheter (i.e., central venous/blood lines, dialysis catheters, nephrostomy tubes, and lumbar punctures); other; and unknown/not stated (unknown).

### Urine collection

Human urine from at least two healthy female donors between the ages of 20 to 35 were collected and pooled. Donors did not have a history of kidney disease, diabetes, or recent antibiotic treatment. Urine was sterilized with a 0.22-μm filter (VWR 29186-212), pH was normalized to 6.0 to 6.5, and urine was used immediately for the assays.

### Fungal cultures

All strains used in this study are listed in Table S1. All strains were cultured at 37°C with aeration in 5 ml of YPD [yeast extract (10 g/liter; VWR J850-500G), peptone (20 g/liter; VWR J636-500G), and dextrose (20 g/liter; VWR BDH9230-500G)] broth. For *in vivo* mouse experiments, *C. auris* strains were grown static overnight in 10 ml of YPD.

### Biofilm formation assays

Biofilm formation assays were performed in 96-well flat-bottomed plates (VWR, 10861-562) coated with 100 μL of Fg (150 μg/mL) incubated overnight at 4°C. The various strains were grown as described above, and the inoculum was normalized to ∼1 × 10^6^ CFUs/ml. Cultures were then diluted (1:100) into human urine supplemented with 10% heat inactivated human serum and incubated in the wells of the 96-well plate at 37°C for 48 hours while static.

### Crystal violet staining

Following biofilm formation on Fg-coated microplate, the supernatant was removed, and the plate was incubated in 200 μL of 0.5% crystal violet for 15 min. Crystal violet stain was removed, and the plate was washed with water to remove the remaining stain. Plates were dried and then incubated with 200 μL of 33% acetic acid for 15 min. In another plate, 100 μL of the acetic acid solution was transferred, and absorbance values were measured via a plate spectrophotometer at 595 nm (Molecular Devices SpectraMax ABS). Values were normalized to AR0382.

### Fibrin(ogen) biofilm formation and visualization

No. 0 cover glass glass-bottom 35-mm petri dish with a 14-mm microwell (MatTek, P35G-0-14-C) was used. For fibrin fiber/nets formation, Fg and thrombin (Sigma-Aldrich, T6884-250UN) were thawed at 37°C. A total of 100 μL of Fg (1 mg/ml) in PBS was added into the microwell glass bottom, and then 10 μL of thrombin (2 U/mL) was added to polymerize Fg into fibrin. Dishes were incubated at 37°C for 1 hour and kept overnight at 4°C.

*C. auris* strains were grown as described above, and the inoculum was normalized to ∼1 × 10^6^ CFUs/ml in PBS. These cultures were then diluted (1:100) into human urine, added into fibrin-coated dishes, and incubated at 37°C for 48 hours under static conditions. After incubation, dishes were then washed three times with 1× PBS to remove unbound fungi, and then dishes were fixed with 10% neutralizing formalin solution for 20 min and washed with 1× PBS three times. Dishes were blocked with blocking solution [1% BSA and 0.3% Triton X-100 (Acros Organics, 21568-2500) in 1× PBS] for 30 min at room temperature. Dishes were incubated with rabbit anti-*Candida* and goat anti-Fg antibodies for 2 hours followed by three washes with PBS-T. Then, dishes were incubated for 1 hour with Alexa Fluor 594–labeled donkey anti-rabbit secondary antibody and Alexa Fluor 488–labeled donkey anti-goat antibodies, followed by three washes with PBS-T. Biofilms were visualized with a Zeiss inverted light microscope, and images were taken at 40x magnification. Zen Pro and Fiji-ImageJ software were used to analyze the images.

### *In vivo* mouse infection models

Mice used in this study were ∼6-week-old female WT C57BL/6 mice purchased from the Jackson Laboratory. **For CAUTI Model:** Mice were subjected to transurethral implantation of a silicone catheter and inoculated as previously described^22^. Briefly, mice were anesthetized by inhalation of isoflurane and implanted with a 6-mm-long silicone catheter (UM catheters; Braintree Scientific, SIL 025). Silicone catheters (LIS) were modified as described in^18^ by modifying medical grade silicone using inert trimethyl-terminated polydimethylsiloxane fluid (silicone oil). Briefly, the oil was absorbed by the silicone tube to create a fully infused silicone tube with a slippery surface. Mice were infected immediately following catheter implantation with 50 μL of ∼1 × 10^6^ CFUs/ml in PBS of one of the fungal strains introduced into the bladder lumen by transurethral inoculation. Mice were euthanized at 1 dpi by cervical dislocation after anesthesia inhalation, and the catheter, bladder, kidneys, spleen, and heart were aseptically harvested. Bladders were weighed and then homogenized for CFUs or fixed in 10% formalin overnight before being paraffin-embedded, sectioned, stained, and imaged. The other organs were homogenized, and catheters were cut into small pieces before sonication for fungal CFU enumeration. **For uUTI Model:** Mice were subjected to transurethral inoculation after being anesthetized by inhalation of isoflurane as previously described^22^. Mice were inoculated with 50 μL of ∼1 × 10^6^ CFUs/ml in PBS of one of the fungal strains. Mice were euthanized at 1 dpi and organs were processed as described in the CAUTI model.

### Immunohistochemistry of mouse bladders

Mouse bladders were fixed in 10% formalin overnight, before being processed for sectioning and staining as previously described^54^. Briefly, bladder sections were deparaffinized, rehydrated, and rinsed with water. Antigen retrieval was accomplished by boiling the samples in Na-citrate, washing in tap water, and then incubating in 1× PBS three times. Sections were then blocked [1% BSA and 0.3% Triton X-100 (Acros Organics, 21568-2500) in 1× PBS], washed in 1× PBS, and incubated with appropriate primary antibodies [Goat anti-Fg, Rabbit anti-Candida, Rat anti-Ly6G] diluted in blocking buffer overnight at 4°C. Next, sections were washed with 1× PBS, incubated with secondary antibodies for 2 hours at room temperature, and washed once more in 1× PBS before Hoechst dye staining. Secondary antibodies for immunohistochemistry were Alexa Fluor 488 donkey anti-goat, Alexa Fluor 550 donkey anti-rabbit, and Alexa Fluor 650 donkey anti-rat. All imaging was done using a Zeiss inverted light microscope. Zen Pro and ImageJ software were used to analyze the images.

### *In vitro* Catheter Biofilms

Disks of UM-silicone (Nalgene 50 silicone tubing, Brand Products) or LIS were cut using an 8 mm leather hole punch. LIS disks were stored in filter sterilized silicone oil at RT. Disks were skewered onto needles (BD) to hold them in place and put in 5 mL glass tubes (Thermo Scientific) and UV sterilized for >30 min prior to use. 500 µL of 150 µg/ml Fg was added to each disk in glass tubes, sealed, and incubated overnight at 4°C. *C. auris* strains were grown as described above, and the inoculum was normalized to ∼1 × 10^6^ CFUs/mL in PBS. These cultures were then diluted (1:100) into human urine, added glass tubes with catheters, and incubated at 37°C for 48 hours under static conditions.

After 48 hours, a sample of the supernatant (urine) was taken to quantify fungal burden in the supernatant by CFUs. The remaining urine was removed, and the catheter piece was washed three times with 1x PBS. Washed catheter pieces were placed in 1 mL of 1xPBS and sonicated for 30 minutes. A sample from the PBS was taken to quantify biofilm CFUs from the catheter.

### Cytokine Analysis

Bladder samples from mice infected (UTI), catheterized and infected with *C. auris* B11103, AR1105, or Chicago4 (CAUTI), or catheterized and mock infected with PBS for 24h were frozen at −80 °C until time of assay. Blood serum samples from mice catheterized and infected with *C. auris* B11103, AR1105, or Chicago4 (CAUTI), or catheterized and mock infected with PBS for 24h were frozen at −80 °C until time of assay. Before cytokine analysis, bladder homogenates thawed on ice and were microcentrifuged at 11,000 × g for 10 min, and supernatants transferred to a new tube. Samples were analyzed using a Bio-Plex Multi-Plex Assay Kit from Bio-Rad Laboratories (Bio-Plex Pro Mouse Cytokine 23-plex Assay #M60009RDPD) following the manufacturer’s protocols. Fold change over naïve bladder or blood serum was calculated for each cytokine.

### Statistical analysis and reproducibility

Data from at least three experiments were pooled for each assay. Two-tailed Mann-Whitney U tests were performed with GraphPad Prism 5 software (GraphPad Software, San Diego, CA) for all comparisons of biofilms, bladder weights, and mouse models. Values represent means ± SEM derived from at least three independent experiments (*P < 0.05; **P < 0.005; ***P < 0.0005; ****P < 0.0001; and ns, difference not significant). For cytokine heatmaps, values represent the median fold change over naïve sample from at least four independent mouse samples. Principal Component Analyses were performed and the PC scores of PC1 and PC2 were plotted for both cytokine (variable) and strain and infection type (loadings).

## Acknowledgments

We thank members of the laboratory of ALFM for helpful suggestions and insightful comments. We give special thanks to Sara Cole, Sarah Chapman, and the ND Integrated Imaging Facility for tissue processing and support during imaging. Funding: This work was supported by institutional funds from the University of Notre Dame (ALFM), and by grants from the Good Venture Foundation (Open Philanthropy, now Coefficient Giving) (to ALFM), from the National Institutes of Health R01DK128805 (to ALFM, AAL, HA, and CH), R01AI177875 (to FHST), and from the Arthur J. Schmitt Leadership Fellowship (to AAL). Additionally, we want to thank Dr. Teresa O’Meara for sharing the Chicago strains with us.

## Author Contributions

ALFM and FHST conceived and supervised the research. AAL, HA, FHST, and ALFM conducted experiments and sample and data analyses. AAL and ALFM wrote the manuscript. ALFM, AAL, FHST, HA, and CH reviewed and edited the final manuscript.

